# The human TRPA1 intrinsic cold and heat sensitivity involves separate channel structures beyond the N-ARD domain

**DOI:** 10.1101/2021.07.31.454589

**Authors:** Lavanya Moparthi, Viktor Sinica, Milos Filipovic, Viktorie Vlachova, Peter M. Zygmunt

**Affiliations:** Wallenberg Centre for Molecular Medicine, Linköping University, SE-581 83 Linköping, Sweden; Department of Biomedical and Clinical Sciences (BKV), Faculty of Health Sciences, Linköping University, SE-581 83 Linköping, Sweden; Department of Cellular Neurophysiology, Institute of Physiology of the Czech Academy of Sciences, 142 20 Prague, Czech Republic; Leibniz-Institut für Analytische Wissenschaften-ISAS-e.V. Bunsen-Kirchhoff-Straße 11, 44139 Dortmund, Germany; Department of Clinical Sciences Malmö, Lund University, SE-214 28 Malmö, Sweden

**Keywords:** Temperature sensing, Cold sensing, Heat sensing, Redox sensitivity, TRP channel, TRPA1, Pain

## Abstract

The human TRPA1 (hTRPA1) is an intrinsic thermosensitive ion channel responding to both cold and heat, depending on the redox environment. Here, we have studied purified hTRPA1 truncated proteins to gain further insight into the temperature gating of hTRPA1. We found in patch-clamp bilayer recordings that Δ1-688 hTRPA1, without the N-terminal ankyrin repeat domain (N-ARD), was more sensitive to cold and heat, whereas Δ1-854 hTRPA1 that is also lacking the S1-S4 voltage sensing-like domain (VSLD) gained sensitivity to cold but lost its heat sensitivity. The thiol reducing agent TCEP abolished the temperature sensitivity of both Δ1-688 hTRPA1 and Δ1-854 hTRPA1. Cold and heat activity of Δ1-688 hTRPA1 and Δ1-854 hTRPA1 were associated with different structural conformational changes as revealed by intrinsic tryptophan fluorescence measurements. Heat evoked major structural rearrangement of the VSLD as well as the C-terminus domain distal to the transmembrane pore domain S5-S6 (CTD), whereas cold only caused minor conformational changes. As shown for Δ1-854 hTRPA1, a sudden drop in tryptophan fluorescence occurred within 25-20°C indicating a transition between heat and cold conformations of the CTD, and thus it is proposed that the CTD contains a bidirectional temperature switch priming hTRPA1 for either cold or heat. In whole-cell patch clamp electrophysiology experiments, replacement of the cysteines 865, 1021 and 1025 with alanine modulated the cold sensitivity of hTRPA1 when heterologously expressed in HEK293T cells. It is proposed that the hTRPA1 CTD harbors cold and heat sensitive domains allosterically coupled to the S5-S6 pore region and the VSLD, respectively.

## 1. Introduction

Several Transient Receptor Potential (TRP) channels are involved in thermosensation allowing organisms to sense a range of temperatures between noxious cold and noxious heat (1-5). Some of these thermoTRPs (TRPV1, TRPV3, TRPM8 and TRPA1) display pronounced inherent temperature sensitivity (6-10), of which the human TRPA1 (hTRPA1) responds to both cold and heat (11). This bidirectional thermosensitivity of hTRPA1, and the findings that chimeras of rat TRPM8 and TRPV1 as well as single point mutations of the mouse TRPA1 caused opposite temperature responses (12, 13) may support the idea that each thermoTRP is both cold and heat sensitive (14). A dual heat and cold sensitivity could involve a single or several specific cold and heat sensing domains that undergo protein denaturation or otherwise are allosterically coupled to the channel gate (14-17). We have reasoned that depending on the cellular environment including the channel redox state, thermoTRPs can adopt various conformations of which some are sensitive to cold and others to heat (7, 11). We have previously shown that hTRPA1 and *Anopheles gambiae* TRPA1 (AgTRPA1) are intrinsically cold and heat activated, respectively, with and without the N-terminal ankyrin repeat domain (N-ARD) and part of the pre-S1 region (7, 8). Furthermore, we found that hTRPA1 is also heat activated and its cold and heat sensitivity most likely involve different protein conformations and possibly separate cold and heat sensors (7, 11, 18).

In this study, we have further investigated the intrinsic cold and heat properties of hTRPA1 beyond the N-ARD, and also with regard to the channel redox state. We found that the heat sensitivity is intact in purified Δ1-688 hTRPA1 (without the N-ARD) but lost in Δ1-854 hTRPA1 (without the N-ARD and S1-S4 transmembrane domain), which displayed an increased cold sensitivity. Different lipid bilayer-independent conformational changes of purified Δ1-688 hTRPA1 and Δ1-854 hTRPA1 caused by cold and heat were disclosed by measuring their intrinsic tryptophan fluorescence. Furthermore, cold and heat responses of purified Δ1-688 hTRPA1 and Δ1-854 hTRPA1 were lost in the presence of the thiol reducing agent TCEP, and cold responses of hTRPA1 subjected to single point mutations of the cysteines 856, 1021 and 1025 were affected when expressed in HEK293T cells. It is suggested that heat sensitivity is dependent on the voltage sensing-like S1-S4 transmembrane domain (VSLD) and the C-terminus domain distal to the transmembrane pore domain S5-S6 (CTD), whereas the cold sensitivity is dependent on the S5-S6 pore region and CTD.

## 2. Materials and Methods

### 2.1. Recombinant protein expression and purification

The expression and purification of Δ1-688 hTRPA1 was performed as described previously (7). The gene sequence of hTRPA1 corresponding to Δ1-854 was optimised for expression in insect cells (GenScript Biotech, Piscataway, NJ, USA). The optimized gene fragment was cloned into the pTriEx-3 baculovirus donor plasmid. The amplification of virus and expression of Δ1-854 hTRPA1 in Hi-5 cells were performed at the Protein Production Platform (LP3, Lund University, Sweden). The Hi-5 cell pellets were lysed using Dounce homogenizer in hypotonic buffer (10 mM HEPES, 36.5 mM sucrose, pH 7.4) supplemented with 0.1% Triton X-100 and a protease inhibitor cocktail. Unbroken cells and cell debris were removed by a short low speed centrifugation of 1500 x g for 5 min. Membranes were collected by ultracentrifugation in a 50.2 Ti rotor (Beckman Instruments) at 45 000 rpm for 1 h. The resulting membrane pellet was solubilized in a membrane storage buffer (20 mM HEPES, 10 % glycerol, 50 mM NaCl, pH 7.8) supplemented with 2% Fos-Choline-14. Detergent insoluble membranes were removed by ultracentrifugation 40,000 rpm at 4 °C for 1 h and the supernatant was incubated with pre-equilibrated Ni-NTA agarose resin for 2 h, at cold temperature. The column was washed with 10 column volumes of buffer containing 20 mM HEPES, 10 % glycerol, 300 mM NaCl, pH 7.8, 0.014 % Fos-Choline-14 and 50 mM imidazole. Δ1-854 hTRPA1 proteins were eluted with the same buffer containing 300 mM instead of 50 mM of imidazole. The protein concentration was determined by measuring absorbance at 280 nm using NanoDrop spectrophotometer. The purity of recombinant proteins was checked by SDS/PAGE followed by Coomassie Blue R-250 staining.

### 2.2. Planar lipid bilayer patch-clamp electrophysiology

These experiments were performed as previously described in detail (7) and are briefly described as follows. Detergent purified hTRPA1 were reconstituted either into preformed planar bilayers or giant unilamellar vesicles (GUVs), composed of 1,2-diphytanoyl-sn-glycero-3-phosphocholine (Avanti Polar Lipids) and cholesterol (Sigma-Aldrich) in a 9:1 ratio and produced by using the Vesicle Prep Pro Station (Nanion Technologies, Germany). Planar lipid bilayers were formed by pipetting 5 µl of either empty GUVs or protein reconstituted into GUVs on patch-clamp chips (1-2 µm, 3.5-5 MΩ resistance) which were mounted on a recording chamber. Following giga Ohm seal formation, single-channel activity was recorded using the Port-a-Patch (Nanion Technologies, Germany) at a holding potential (Vh) of +60 mV in a symmetrical K^+^ solution adjusted to pH 7.2 with KOH and containing (in mM): 50 KCl, 10 NaCl, 60 KF, 20 EGTA, and 10 HEPES. The patch-clamp experiments were performed at various temperatures, for which the Port-a-Patch was equipped with an external perfusion system (Nanion Technologies) and an SC-20 dual in-line solution cooler/heater connected to a temperature controlled (CL-100) liquid cooling system (Warner Instruments). Signals were acquired with an EPC 10 amplifier (HEKA) and the data acquisition software Patchmaster (HEKA) at a sampling rate of 50 kHz. The recorded data were digitally filtered at 3 kHz. Electrophysiological data were analysed using Clampfit 9 (Molecular Devices) and Igor Pro (Wave Metrics software). Data were filtered at 1000 and 500 Hz low-pass Gaussian filter for analysis and traces, respectively. The single-channel mean conductance (Gs) was obtained from a Gaussian fit of all-points amplitude histograms. The single-channel mean open probability (Po) was calculated from time constant values, which were obtained from exponential standard fits of dwell time histograms. The Q_10_ for Po was obtained from the following equation (19, 20):

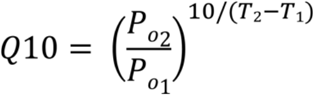

### 2.3. Whole-cell patch-clamp electrophysiology

#### 2.3.1. Cell culture, constructs and transfection

Human embryonic kidney 293T (HEK293T; ATCC, Manassas, VA, USA) cells were cultured in Opti-MEM I media (Invitrogen, Carlsbad, CA, USA) supplemented with 5% fetal bovine serum. The magnet-assisted transfection (IBA GmbH, Gottingen, Germany) technique was used to transiently co-transfect the cells in a 15.6 mm well on a 24-well plate coated with poly-L-lysine and collagen (Sigma-Aldrich, Prague, Czech Republic) with 200 ng of GFP plasmid (TaKaRa, Shiga, Japan) and with 300 ng of cDNA plasmid encoding wild-type or mutant human TRPA1 (pCMV6-XL4 vector, OriGene Technologies, Rockville, MD, USA). The cells were used 24–48 h after transfection. At least three independent transfections were used for each experimental group. The wild-type channel was regularly tested in the same batch as the mutants. Coding sequences of hTRPA1 mutants were synthesized and sub cloned into the vector pcDNA3.1(+) by GenScript Biotech (Piscataway, NJ, USA).

#### 2.3.2. Whole-cell electrophysiology and temperature stimulation

Whole-cell membrane currents were filtered at 2 kHz using the low-pass Bessel filter of the Axopatch 200B amplifier and digitized (5-10 kHz) using a Digidata 1440 unit and pCLAMP 10 software (Molecular Devices, San Jose, CA, USA). Patch electrodes were pulled from borosilicate glass and heat-polished to a final resistance between 3 and 5 MΩ. Series resistance was compensated by at least 60%. Only one recording was performed on any one coverslip of cells to ensure that recordings were made from cells not previously exposed to temperature. Extracellular bath solutions were Ca^2+^-free and contained: 140 mM NaCl, 5 mM KCl, 2 mM MgCl_2_, 5 mM EGTA, 10 mM HEPES, 10 mM glucose, pH 7.4 was adjusted by TMA-OH. Intracellular solution contained 140 mM KCl, 5 mM EGTA, 2 mM MgCl_2_, 10 mM HEPES, adjusted with KOH to pH 7.4. The *I-V* relationships were recorded using 200-ms ramps ranging from -100 mV to +100 mV, holding potential 0 mV. Voltage ramp protocol (1 V/s; shown in the inset of Fig. 6A) was applied repeatedly each 1 s. A system for fast cooling and heating of solutions superfusing isolated cells under patch-clamp conditions was used as described previously (21).

### 2.4. Intrinsic tryptophan fluorescence assay

Conformational changes in Δ1-688 hTRPA1 and Δ1-854 hTRPA1 were recorded on FP-8200 spectrofluorometer (Jasco, Germany) with thermostat unit, as previously described for hTRPA1 (11). Protein was incubated at each desired temperature for 15 min and emission spectra recorded using excitation wavelength of 280 nm. Cold and heat experiments were done in separate experiments and for better presentation the spectra are always normalized to the starting spectrum at room temperature. To keep the same oxygen levels in all samples and to prevent the spontaneous protein aggregation/precipitation, the measurements were performed in the sealed anaerobic cuvettes with constant stirring.

### 2.5. Statistics

SigmaPlot 10 (Systat Software Inc., San Jose, USA) and GraphPad Prism 9.1. (GraphPad Software, La Jolla, CA) were used for statistical analysis and drawing of graphs. The level of statistical significance was set at *P* < 0.05. One-way ANOVA and Dunnett’s post hoc comparison tests were used for analysis of statistical significance. Data are presented as the mean ± SEM; *n* indicates the number of separate experiments examined.

## 3. Results

### 3.1. Effect of temperature on purified Δ1-688 hTRPA1 and Δ1-854 hTRPA1 channel activity

The purified Δ1-688 hTRPA1, reconstituted into artificial planar lipid bilayers, responded with pronounced channel activity to temperatures above 25°C (Fig. 1A, Table 1). A near maximum single-channel open probability value was reached already at 30°C with a calculated Q_10_ value of 29. In agreement with previous studies (7, 11), Δ1-688 hTRPA1 was activated by cold (Fig. 1A, Table 1). The purified Δ1-854 hTRPA1, reconstituted into artificial planar lipid bilayers, also responded with pronounced channel activity but to temperatures below 20°C (Fig. 2A, Table 2). Within the applied temperature interval of 20-15°C, the single-channel open probability value reached a maximum close to 1 (Fig. 2C) with a calculated Q_10_ value of 1000. Although, the heat sensitivity was lost, Δ1-854 hTRPA1 was active with similar single-channel open probability values within a temperature interval of 22-40°C (Fig. 2).

**Table 1.** Single-channel open probability and conductance values for Δ1-688 hTRPA1.

| Stimuli | Voltage | Open probability | n | Conductance (pS) | n |
| --- | --- | --- | --- | --- | --- |
| | (mV) | (mean ± SEM) | | (mean ± SEM) | |
| 40 °C | +60 | 0.73 ± 0.09 | 3 | 54 ± 12 | 3 |
| 35 °C | +60 | 0.64 ± 0.04 | 9 | 27 ± 4 | 9 |
| 30 °C | +60 | 0.58 ± 0.04 | 18 | 43 ± 4 | 18 |
| 25 °C | +60 | 0.05 ± 0.05 | 8 | - | 8 |
| 22 °C | +60 | 0.00 ± 0.00 | 4 | - | 4 |
| 20 °C* | +60 | 0.02 ± 0.02 | 3 | 68 ± 17 | 3 |
| 17 °C* | +60 | 0.22 ± 0.04 | 4 | 54 ± 10 | 4 |
| 15 °C | +60 | 0.57 ± 0.06 | 5 | 23 ± 2 | 5 |
| 10 °C* | +60 | 0.93 ± 0.02 | 4 | 15 ± 3 | 4 |

\* data published previously (ref. 7).

**Table 2.** Single-channel open probability and conductance values for Δ1-854 hTRPA1.

| Stimuli | Voltage | Open probability | n | Conductance (pS) | n |
| --- | --- | --- | --- | --- | --- |
| | (mV) | (mean ± SEM) | | (mean ± SEM) | |
| 40 °C | +60 | 0.24 ± 0.07 | 5 | 31 ± 5 | 5 |
| 35 °C | +60 | 0.17 ± 0.08 | 3 | 54 ± 16 | 3 |
| 30 °C | +60 | 0.20 ± 0.04 | 12 | 33 ± 4 | 12 |
| 25 °C | +60 | 0.25 ± 0.12 | 3 | 33 ± 7 | 3 |
| 22 °C | +60 | 0.22 ± 0.02 | 3 | 23 ± 2 | 3 |
| 20 °C | +60 | 0.03 ± 0.02 | 5 | 17 ± 0 | 2 |
| 17 °C | +60 | 0.33 ± 0.04 | 6 | 34 ± 6 | 6 |
| 15 °C | +60 | 0.92 ± 0.03 | 9 | 33 ± 4 | 9 |

**Fig. 1.**
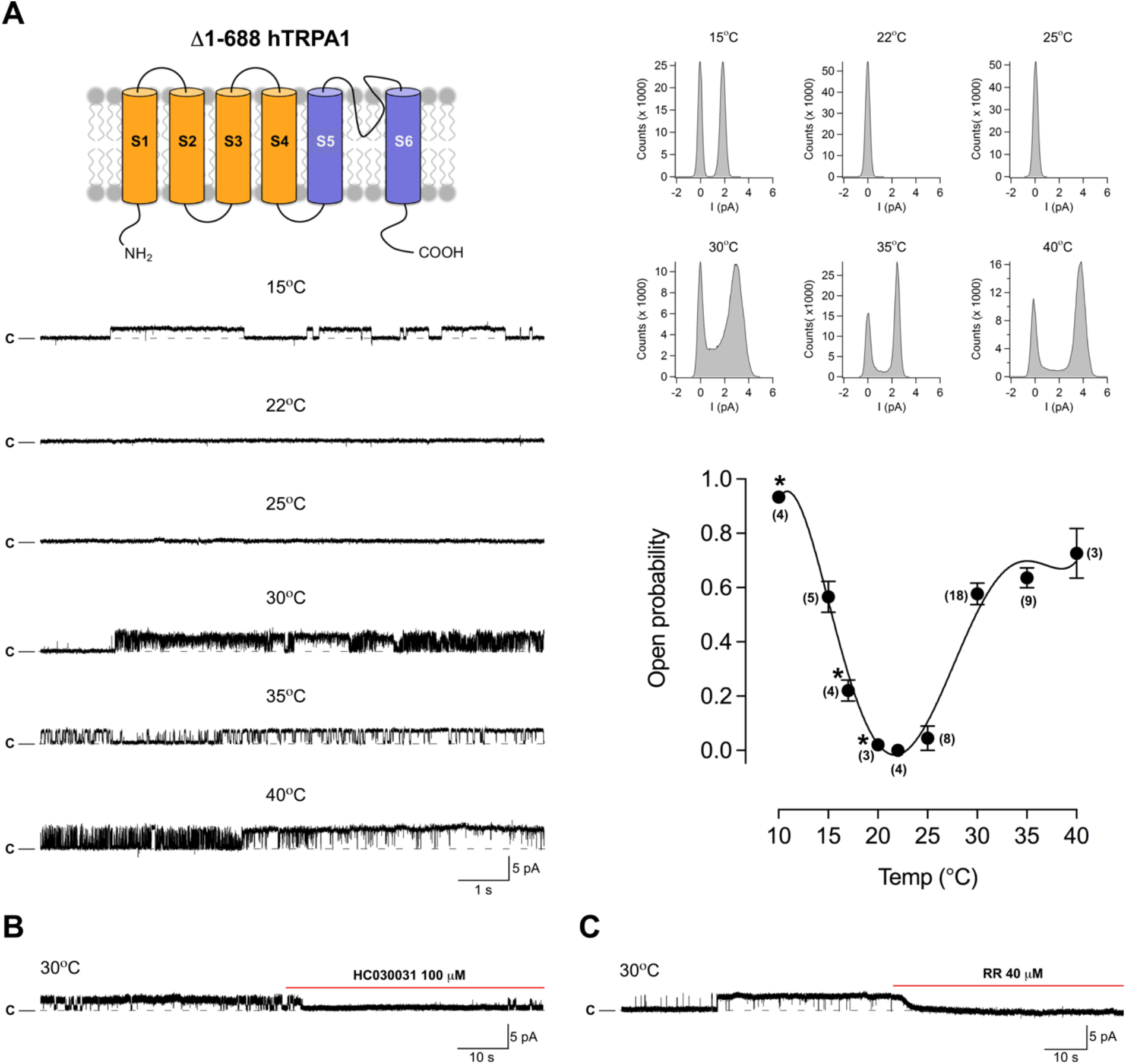
The purified human TRPA1 without its N-terminal ankyrin repeat domain (Δ1-688 hTRPA1) is both cold- and heat sensitive. Purified Δ1-688 hTRPA1 was reconstituted into planar lipid bilayers and single-channel currents were recorded with the patch-clamp technique in a symmetrical K^+^ solution at a holding potential of +60 mV. (**A**) As shown by representative traces and corresponding histograms, exposure to various temperatures evoked outward single-channel currents. The graph shows single-channel mean open probability values as a function of different temperatures. Data are represented as the mean ± SEM of independent experiments (shown within parentheses). Data points marked with asterisk were published previously (ref. 7). (**B** and **C**) The selective TRPA1 antagonist HC030031 and the non-selective TRP channel pore blocker ruthenium red (RR) inhibited heat responses (each n = 3). Black dotted line shows zero channel current level (c indicates the close channel state) and upward deflections are channel openings. Q_10_ cold = 56 (ref. 7); heat (20°C to 30°C) = 29.

**Fig. 2.**
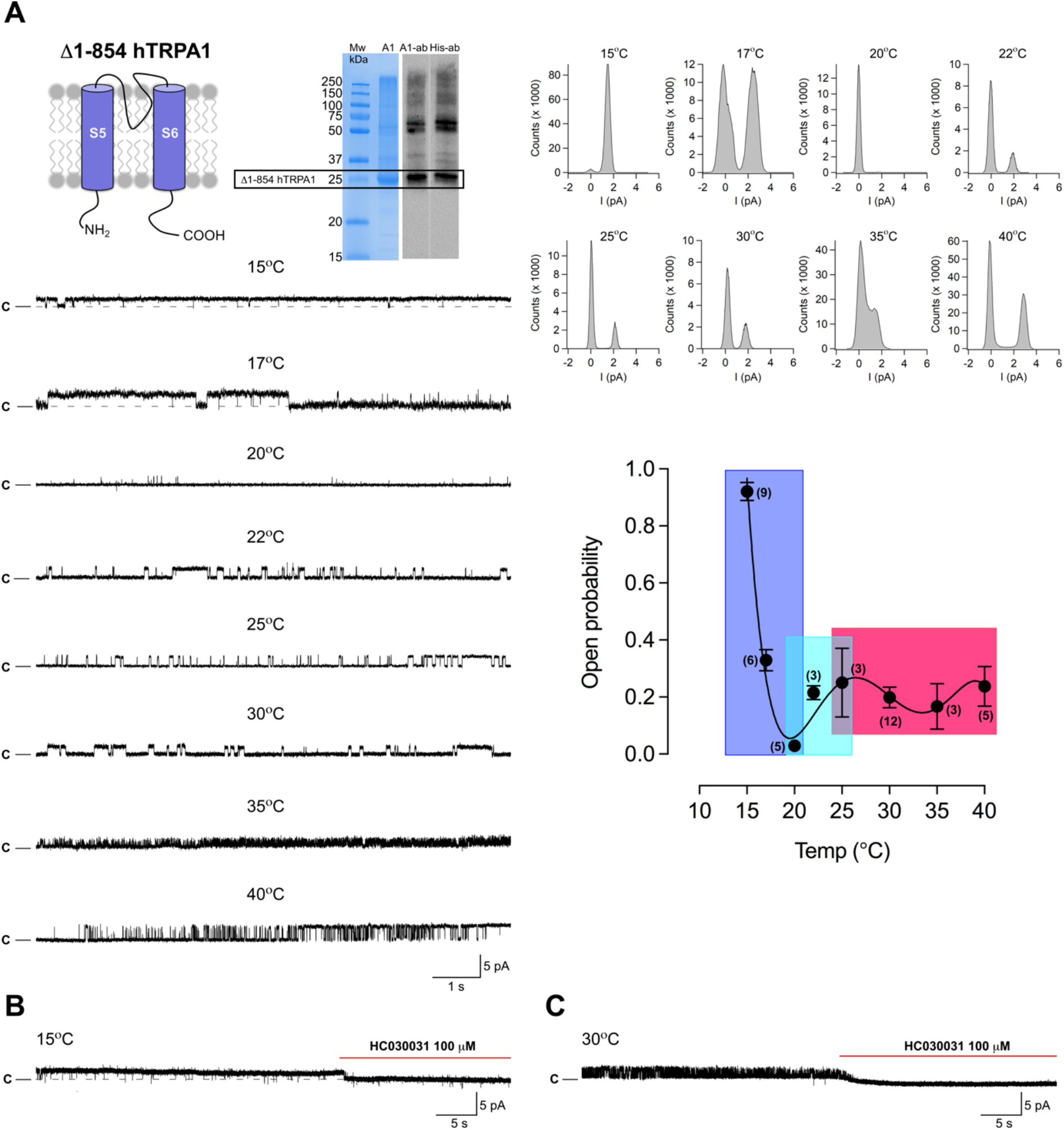
The purified human TRPA1 without its N-terminal ankyrin repeat domain and transmembrane segments S1-S4 (Δ1-854 hTRPA1) is cold- but not heat sensitive. Purified Δ1-854 hTRPA1 was reconstituted into planar lipid bilayers and single-channel currents were recorded with the patch-clamp technique in a symmetrical K^+^ solution at a holding potential of +60 mV. (**A**) Affinity-purified Δ1-854 hTRPA1 is visualized by Coomassie staining (left gel), and by Western blotting (right gel) using either hTRPA1 antibody (left lane) or tetrahistidine antibody (right lane). As shown by representative traces and corresponding histograms, exposure to various temperatures evoked outward single-channel currents. The graph shows single-channel mean open probability values as a function of different temperatures. Data are represented as the mean ± SEM of independent experiments (shown within parentheses). Coloured areas relate to the Δ1-854 hTRPA1 intrinsic tryptophan fluorescence measured within the same temperature interval (Fig. 3). (**B** and **C**) The selective TRPA1 antagonist HC030031 inhibited both cold and heat responses (each n = 4). Black dotted line shows zero channel current level (c indicates the close channel state) and upward deflections are channel openings. Q_10_ cold (20°C to 15°C) = 1000.

The activity of Δ1-688 hTRPA1 and Δ1-854 hTRPA1 was abolished by the TRPA1 inhibitors HC030031 and ruthenium red (Fig. 1B,C and 2B,C), at concentrations that abolished purified hTRPA1 and Δ1-688 hTRPA1 single-channel activity evoked by chemical ligands, temperature and mechanical stimuli (7, 11, 22-24). Furthermore, no channel activity was observed in membranes, without the purified hTRPA1 proteins in the presence of the detergent Fos-Choline-14, when exposed to the temperatures used in the present study (7, 11, 22-24).

### 3.2. Effect of temperature on purified Δ1-688 hTRPA1 and Δ1-854 hTRPA1 intrinsic tryptophan fluorescence activity

To study lipid bilayer-independent conformational changes in response to cold and heat, we exposed purified detergent solubilized Δ1-688 hTRPA1 and Δ1-854 hTRPA1 to various temperatures and measured the intrinsic tryptophan fluorescence signal (Fig. 3). The maximum fluorescence signal was larger in Δ1-688 hTRPA1 compared to Δ1-854 hTRPA1, most likely because of more tryptophans in Δ1-688 hTRPA1 (Fig. 4). Heat almost completely extinguished the Δ1-688 hTRPA1 fluorescence signal and caused substantial quenching of the Δ1-854 hTRPA1 fluorescence signal (Fig. 3). When lowering the temperature from 25°C to 19°C a sudden major drop of the Δ1-854 hTRPA1 fluorescence signal appeared, after which only a minor change of the protein fluorescence signal was observed with further cooling (Fig. 3). A small gradual decrease in tryptophan fluorescence was also observed for Δ1-688 hTRPA1 when lowering the temperature from 22°C (Fig. 3). In the presence of carvacrol, cold caused a larger quenching of Δ1-688 hTRPA1 fluorescence (Fig. 3).

**Fig. 3.**
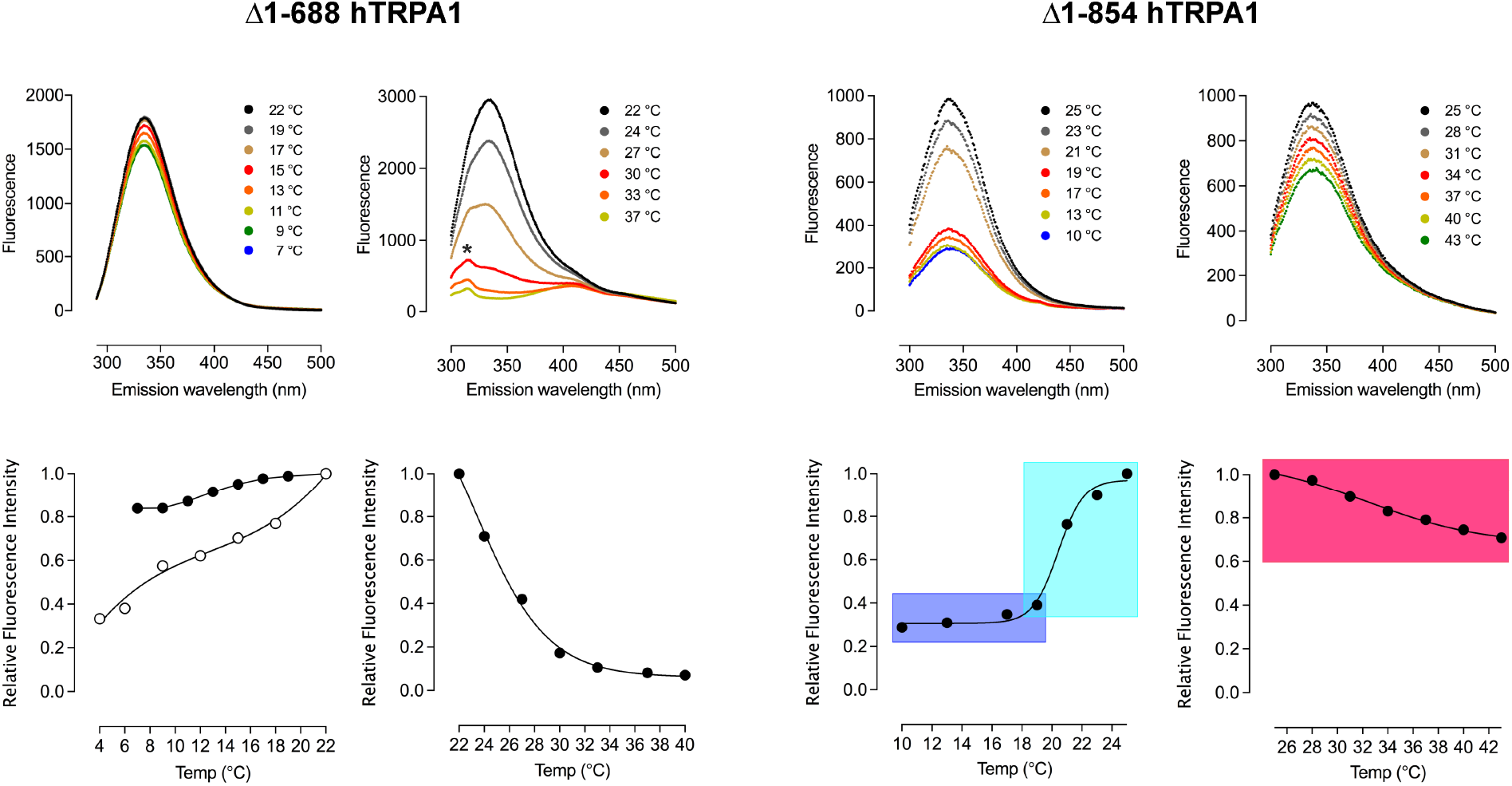
Cold and heat cause lipid bilayer-independent conformational changes of purified Δ1-688 hTRPA1 and Δ1-854 hTRPA1. Spectra show different conformational changes induced by cold or heat. The hump (*) was only observed for Δ1-688 hTRPA1 and above 22°C. Coloured areas relate to the Δ1-854 hTRPA1 channel open probability measured within the same temperature interval (Fig. 2). Shown is also the effect of carvacrol (100 μM, open circles) on Δ1-688 hTRPA1 cold responses. At 22°C, carvacrol itself emitted fluorescence that was subtracted when its effect on cold was determined. The intrinsic tryptophan fluorescence intensity, emitted at 335 nm, for each indicated temperature was related to that of 22°C or 25°C and expressed as Relative Fluorescence Intensity in the graphs, where data are represented as the mean ± SEM of three independent experiments.

**Fig. 4.**
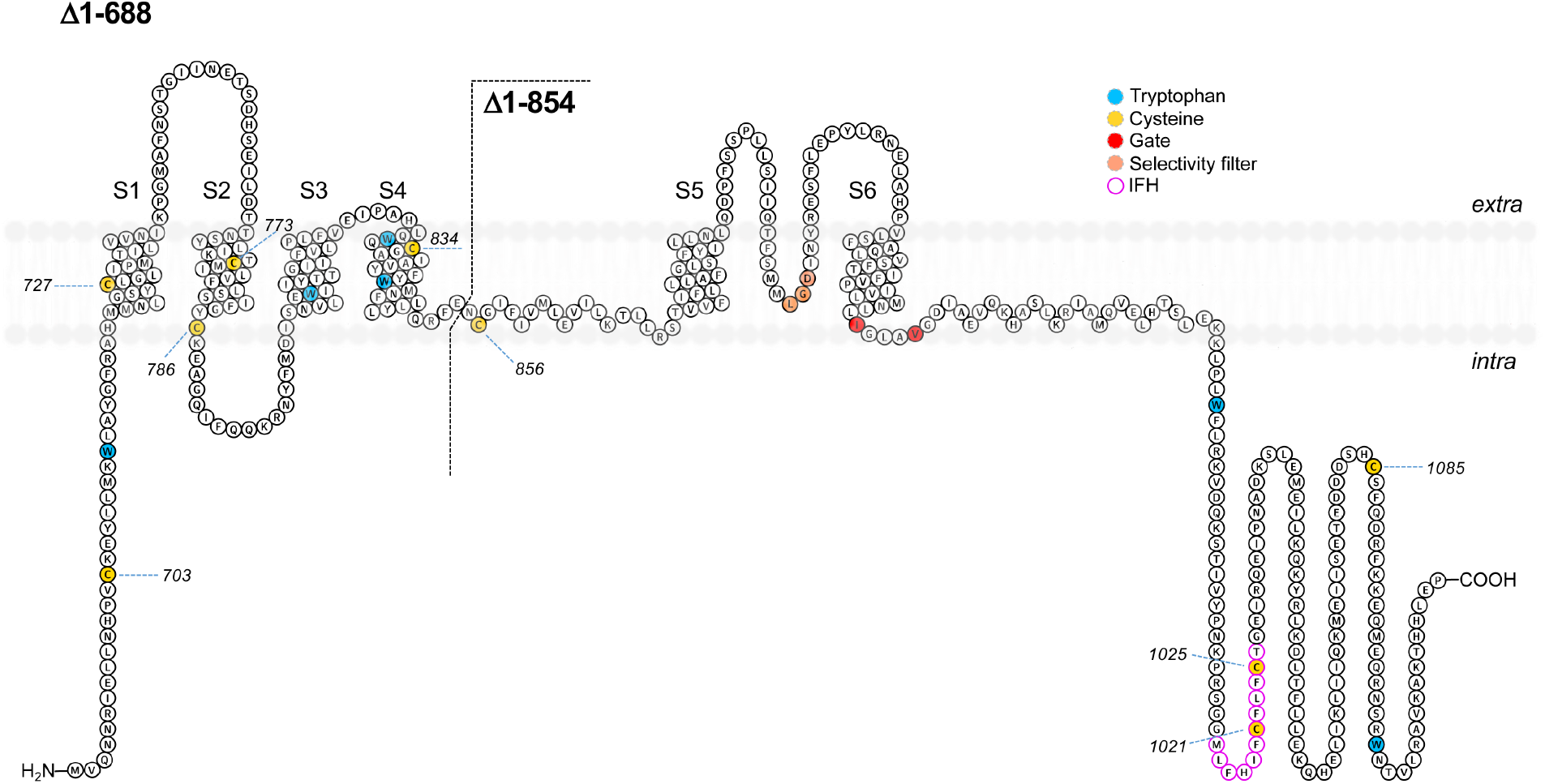
Topology plot of Δ1-688 hTRPA1 and Δ1-854 hTRPA1. Shown is hTRPA1 without its N-ARD and part of the pre-S1 region (Δ1-688 hTRPA1) and its further truncated isoform (Δ1-854 hTRPA1). Highlighted are tryptophans with the capacity to contribute to the intrinsic fluorescence recorded in Fig. 3, and cysteines of which the role of C856, C1021 and C1025 in hTRPA1 cold responses were explored in whole-cell patch clamp studies (Fig. 6). C1021 and C1025 are located within a short helix, termed interfacial helix (IFH), that makes close contact with the voltage sensing-like domain (VSLD) consisting of S1-S4.

### 3.3. Effect of TCEP on purified Δ1-688 hTRPA1 and Δ1-854 hTRPA1 temperature-dependent channel activity

Both Δ1-688 hTRPA1 and Δ1-854 hTRPA1 contain cysteines that may become oxidized and form various intra- and intermolecular disulphide networks affecting channel function properties (Fig. 4). As shown previously, hTRPA1 cold- and heat responses, and Δ1-688 hTRPA1 cold responses were prevented by the thiol reducing agent TCEP (7, 11). As shown in the present study, TCEP at the same concentration (1 mM) also completely prevented heat responses in Δ1-688 hTRPA1 as well as Δ1-854 hTRPA1 cold responses (Fig.5). The effect of TCEP on Δ1-854 hTRPA1 heat responses was not investigated as its heat sensitivity was lost (Fig. 2).

**Fig. 5.**
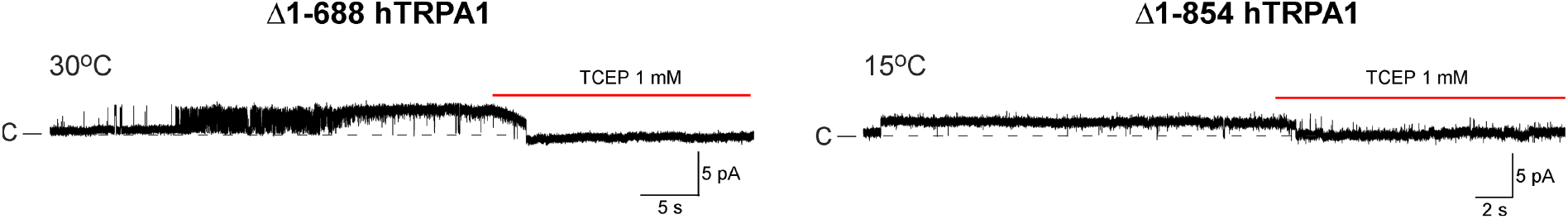
Δ1-688 hTRPA1 and Δ1-854 hTRPA1 cold and heat sensitivity is dependent on the channel redox state. Purified Δ1-688 hTRPA1 was reconstituted into planar lipid bilayers and single-channel currents were recorded with the patch-clamp technique in a symmetrical K^+^ solution at a holding potential of +60 mV. As shown by traces, the thiol reducing agent TCEP eliminated the channel activity of Δ1-688 hTRPA1 evoked by heat (n = 6) and Δ1-854 hTRPA1 evoked by cold (n = 5). Black dotted line shows zero channel current level (c indicates the close channel state) and upward deflections are channel openings.

### 3.4. Whole-cell cold activated currents in HEK293T cells expressing hTRPA1 cysteine mutants

The role of cysteines 856, 1021 and 1025 in voltage-dependent cold responses was examined by measuring whole-cell currents in HEK293T cells expressing three single cysteine mutants (C856A, C1021A, C1025A) and one double cysteine mutant (C1021A/C1025A) of hTRPA1 (Fig. 6). Repeated voltage ramps from –100 mV to +100 mV were applied in control extracellular solution and current-voltage (*I*-*V*) relations were compared at 25 °C and 15 °C (Fig. 6A-D). At 25°C, the C856A and C1025A constructs exhibited a significantly stronger outward rectification (current at -80 mV/current at +80 mV; 0.07 ± 0.02 and 0.06 ± 0.01; n = 11 and 14) compared with wild-type channels (0.16 ± 0.02; n = 26), indicating that the mutations modified the voltage-dependent gating. At -80 mV, cold (15°C) potentiated the currents through wild-type channels by 1.9 ± 0.1-fold and the extent of modulation by cold was significantly different in the C856A and C1025A mutants (Fig. 6E). The C1025A construct was potentiated by cold by 2.7 ± 0.3-fold, whereas the currents through C856A did not exhibit any significant change at -80 mV (1.2 ± 0.1). At positive membrane potentials (+80 mV), cold reduced the outward currents through wild-type hTRPA1 by 0.63 ± 0.02-fold but the currents mediated by C1021A and C1025A were reduced only by 0.50 ± 0.3- and 0.45 ± 0.02-fold (Fig. 6E). Thus, in all three single cysteine mutants tested, the extent of modulation by cold was significantly altered at either negative or positive membrane potential. Interestingly, the double mutant C1021A/C1025A restored hTRPA1 cold responses (Fig. 6D) and exhibited an outward rectification at 25°C not different from wild-type channels (0.22 ± 0.03; P = 0.472; n = 11). Together, these results suggest that the three CTD cysteines are involved in both the temperature- and voltage-dependent gating of hTRPA1.

**Fig. 6.**
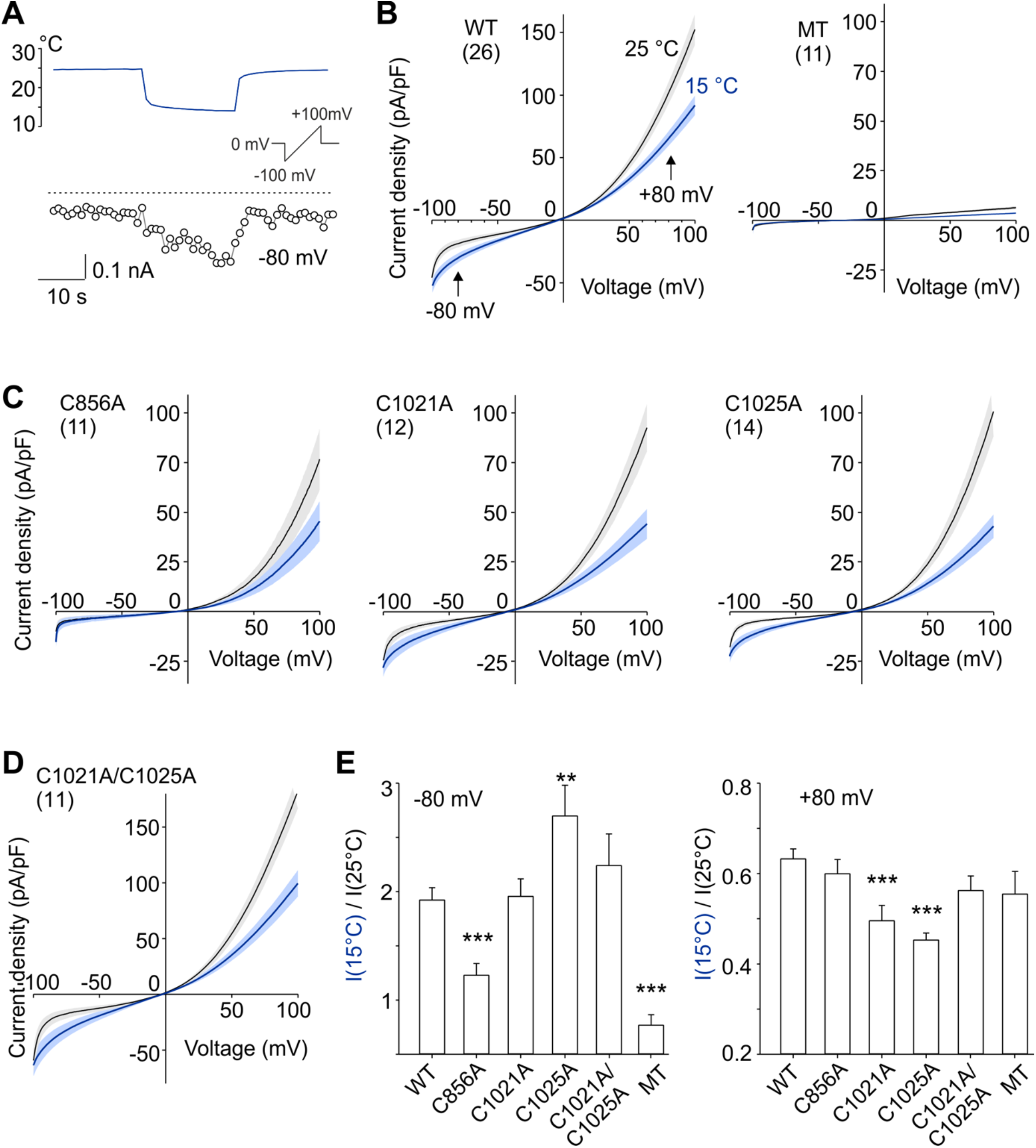
The voltage-dependent activation properties of cysteine mutants of human TRPA1, C856A, C1021A, C1025A and C1021A/C1025A, are differentially modulated by cold. (**A**) Representative time course of whole-cell current amplitudes measured at -80 mV from a HEK293T cell expressing human TRPA1 channels. Voltage ramp protocol (1 V/s; shown in the inset) was applied repeatedly each 1 s. (**B**) Average current-voltage (*I-V*) relations obtained at 25°C (black lines and gray envelopes are the mean ± SEM) and at 15°C (blue lines and lighter envelopes are the mean ± SEM) from wild-type human TRPA1 (WT; left) or mock transfected cells (MT; right). (**C, D**) Average *I-V* relations obtained at 25°C and at 15°C from the indicated cysteine mutants. Number of independent experiments is shown in parentheses for each construct. (**E**) Bar graph summarizing the effects of cold measured at -80 mV and +80 mV (see vertical arrows in **B**). Data are represented as mean ± SEM. **P < 0.01 and ***P < 0.001 indicate statistically significant differences compared to WT using the One-way ANOVA and Dunnett’s post hoc comparison tests.

## 4. Discussion

Several TRP channels are involved in mammalian thermosensation with some overlap in temperature responsiveness (1-5). but only few thermoTRPs have been shown to respond directly to a change in temperature (6, 8-11). In our studies on purified hTRPA1 and AgTRPA1, we found that hTRPA1 responded to cold and AgTRPA1 to heat, and that these intrinsic temperature responses remained in the absence of the N-ARD and part of pre-S1 region (7, 8). We speculated that the redox state and/or other post-translational modifications controlled the hTRPA1 intrinsic temperature sensitivity also allowing hTRPA1 to be activated by heat (11), which would support findings indicating an involvement of TRPA1 in mammalian heat detection *in vivo* (25-30). Indeed, cold- and heat-induced activity of hTRPA1 was found to be dependent on the channel redox state, and both oxidants and ligand activators such as hydrogen peroxide (H_2_O_2_), allyl isothiocyanate (AITC), acrolein and carvacrol modified hTRPA1 temperature activation in intact HEK293 cells and mouse trachea sensory neurons (11). We therefore concluded that the heat sensing property of TRPA1 is conserved in mammalians, in which TRPA1 may contribute to sensing warmth and uncomfortable heat in addition to noxious cold. Similar conclusions regarding TRPA1 as a heat sensor of physiological relevance have been reached in recent publications (18, 31-33).

In this study, we show that hTRPA1 without its N-ARD (Δ1-688 hTRPA1) retains its bidirectional thermosensitive profile but with increased thermosensitivity as indicated by Q_10_ values of 40 (hTRPA1) and 56 (Δ1-688 hTRPA1) for cold, and 6 (hTRPA1) and 29 (Δ1-688 hTRPA1) for heat. Likewise, the heat-sensitivity of Ag-TRPA1(A) without its N-ARD (Δ1-776 AgTRPA1) was estimated higher than for Ag-TRPA1(A) for which a Q_10_ value of 28 could be calculated (8). This suggests a modulatory role of the N-ARD in TRPA1 thermosensation, but may not necessarily be driven by temperature sensitive domains within N-ARD as both calcium responses and mechanosensitivity were different between purified hTRPA1 and Δ1-688 hTRPA1 at 22°C and otherwise identical experimental conditions (23, 24). Further deletion of VSLD in hTRPA1 (Δ1-854 hTRPA1), keeping most of the S4-S5 linker with the redox sensitive C856 and N855, the mutation of which in humans causes stress-induced cold hypersensitivity (34, 35), created a channel with a dramatic increase in cold sensitivity and lost heat sensitivity, the latter indicating heat sensitive structures within VSLD or its uncoupling from a heat sensitive CTD. The increase in cold sensitivity with truncation as demonstrated by Q_10_ values of 40 (hTRPA1), 56 (Δ1-688 hTRPA1) and 1000 (Δ1-854 hTRPA1) suggests that N-ARD and VSLD negatively modulate the cold-induced responses, which may be triggered by thermosensor loci within both S5-S6 and CTD. Interestingly, this “break” of cold responses can be counteracted by both electrophilic (AITC and acrolein) and non-electrophilic (carvacrol) TRPA1 activators, interacting with binding sites within the N-ARD and VSLD (7, 31, 36-38), as shown by their ability to sensitize mammalian TRPA1 cold responses (11, 18, 31, 39). The selective TRPA1 antagonist HC030031 completely prevented cold and heat responses of hTRPA1 and its truncated isoforms in previous and present studies (7, 11) supporting the idea that HC030031 inhibits hTRPA1 by interacting with N855 and the CTD (40).

Because hTRPA1 is an inherent mechanosensitive ion channel gated by force-from-lipids (24), it could be argued that hTRPA1 is indirectly thermosensitive by responding to temperature-dependent bilayer fluidity changes. However, the lipid 1,2-diphytanoyl-sn-glycero-3-phosphocholine form bilayers that are structurally stable within the temperature interval studied here and in previous studies (41), and in which no currents were detected in the absence of hTRPA1 when exposed to the same test temperatures (7, 8, 11). Further evidence of hTRPA1 being intrinsically thermosensitive is provided by lipid bilayer-independent measurements of temperature-dependent hTRPA1 intrinsic tryptophan fluorescence activity.

The intrinsic tryptophan fluorescence spectroscopy measurements, in contrast to patch-clamp recordings, can detect structural channel rearrangements associated with temperature changes. The stronger tryptophan fluorescence signal in Δ1-688 hTRPA1 compared to Δ1-854 hTRPA1 indicates a major contribution from the tryptophans located within S3-S4 and the pre-S1 region that may mask the detection of any CTD conformational changes, signalled by only its two tryptophans, in response to heat and cold. The extensive heat-induced quenching of the Δ1-688 hTRPA1 tryptophan fluorescence, and the characteristic hump on the Δ1-688 hTRPA1 but not Δ1-854 hTRPA1 fluorescence spectra at lower emission wavelengths, suggest that the heat temperature sensor is located within the VSLD. However, structural rearrangements of VSLD could possibly also occur as a result of the heat-induced conformational changes of the CTD as revealed by Δ1-854 hTRPA1. Thus, it could be that heat is properly detected by CTD, but without coupling to the VSLD heat cannot gradually be transformed into gating of the channel, as shown by the lost heat-sensitivity in Δ1-854 hTRPA1. Notably, some conformational change of the pore region must still occur in the absence of VSLD within 25-40°C, since single-channel currents were still recorded, albeit with single-channel open probability independent of temperature. Importantly, the Δ1-854 hTRPA1 channel activity at 30°C was abolished by HC030031 and its single-channel conductance within 25-40°C was not different from that of Δ1-688 hTRPA1 in the present study and hTRPA1 (11), indicating that uncoupling of CTD and VSLD affected only the heat-sensitivity but not the pore function of Δ1-854 hTRPA1. Thus, it could be that a heat sensor is located to either CTD or VSLD, or present in both CTD and VSLD possibly acting in concert to gate hTRPA1. The cold-induced tryptophan fluorescence changes of Δ1-688 hTRPA1 were modest and could be the result of minor rearrangement of the VSLD, sensing cold directly or indirectly by cold-evoked structural changes of S5-S6, which lacks tryptophans. Furthermore, the small cold-evoked structural rearrangement of the CTD, as shown for Δ1-854 hTRPA1 when lowering the temperature from around 20°C, may also take part in Δ1-688, but is not visible because of the strong tryptophan signal from VSLD. Importantly, tryptophan fluorescence measurements of Δ1-854 hTRPA1 revealed a sudden drop in tryptophan fluorescence within 25-20°C indicating a transition between heat and cold conformations of the CTD, and thus it is proposed that the CTD contains a bidirectional temperature switch priming hTRPA1 for either cold or heat. This fits well with the obtained single-channel open probability nadir at 20-22°C observed for hTRPA1 (11) and its truncated isoforms in the present study. Thus, it is possible that cold and heat denaturation of CTD, which in hTRPA1 is in close contact with the pre-S1 region and the S4-S5 linker connecting the VSLD and pore domain, is a key mechanism in the temperature gating of hTRPA1. Interestingly, swapping the CTD between TRPV1 and TRPM8 caused opposite temperature sensitivity (12). Furthermore, it was suggested that a folded-unfolded transition of a specialized temperature-sensitive structure in the CTD of TRPM8 is associated with an increase in the heat molar capacity and required for its gating by cold (16).

As mentioned earlier, electrophiles and carvacrol sensitized the cold-induced hTRPA1 whole-cell currents studied in HEK293T cells (11, 18, 31, 39). Furthermore, the cold-evoked hTRPA1 conformational change as measured by tryptophan fluorescence signaling was much larger in the presence of carvacrol (11). Whereas carvacrol most likely binds within the VSLD (31), electrophilic TRPA1 activators are supposed to trigger responses by interacting with highly reactive cysteines (Cys621, Cys641 and Cys665) in the hTRPA1 N-ARD/pre-S1 region (42-44). However, electrophiles such as N-methyl maleimide (NMM), (E)-2-alkenals, AITC, cinnamaldehyde and *p*-benzoquinone can activate TRPA1 by interacting with other cysteines and lysines outside the N-ARD (7, 38, 44-46). Interestingly, as determined by mass spectrometry, the NMM labelling of C834 in S4 decreased with increasing concentrations of NMM in both hTRPA1 and Δ1-688 hTRPA1, indicating a structural rearrangement of S4 by electrophiles (38) as confirmed by cryo-EM (43, 44). In the present study, we further examined the effect of carvacrol on the intrinsic tryptophan fluorescence activity in Δ1-688 hTRPA1 in response to cooling. In the presence of carvacrol, the tryptophan fluorescence was extensively quenched by cold temperatures which otherwise only produced a minor change in tryptophan fluorescence intensity. Taken together, these findings support a ligand-induced removal of a “cold break” in the VSLD.

The purified hTRPA1 is partially oxidized and thiol modifying agents can either inhibit or potentiate its temperature responses (11). In the present study, we found that Δ1-688 hTRPA1 heat-induced activity was abolished by the thiol reducing agent TCEP, which also inhibited the cold-induced activity of Δ1-688 hTRPA1 as well as both cold and heat responses of hTRPA1 (11). Likewise, TCEP prevented cold-evoked Δ1-854 hTRPA1 channel activity in the present study. Taken together, these findings show that the redox state of hTRPA1 is a critical factor determining its temperature sensitivity. The number of cysteines is substantially reduced with truncation and as a consequence the intra/inter molecular disulphide network is most likely different in the hTRPA1 and its truncated isoforms. However, we reasoned that the cysteines 856, 1021 and 1025 are still important in coordinating CTD coupling to the pore region of Δ1-854 hTRPA1 as well as to the VSLD and pre-S1 region in Δ1-688 hTRPA1 and hTRPA1 (31). We therefore decided to further explore the role of these cysteines in cold-evoked TRPA1 activation by expressing hTRPA1 in HEK293T cells for patch-clamp studies.

Our whole-cell patch-clamp studies on mutant channels show that the cysteines 856, 1021 and 1025 are involved in the cold-dependent activation of hTRPA1. In particular, C856A was not affected at all by cold at negative membrane potentials, whereas the wild-type channels were potentiated by about two-fold. This residue has previously been recognized as one of the main targets of O_2_ in hyperoxia (35), but it is also critical for the gating equilibrium and/or voltage-dependent activation of the channel (31). The important structural role of this residue is clearly supported by recently resolved structures of hTRPA1 in different conformational states (43, 44, 47). C856 is located at the N-terminal portion of the S4-S5 linker, which is directly involved in conformational changes leading to channel opening by covalent agonists. Interestingly, in the closed conformation under the apo conditions (PDB ID: 6V9W), C856 contacts V875 in the fifth transmembrane domain S5 of the neighboring subunit. In contrast, the distance between C856 and V875 is more than 15 Å in a conformation opened by the irreversible electrophilic agonist iodoacetamide (PDB ID: 6V9X). The valine 875 has been identified as a molecular determinant underlying the species-specific differences in cold sensitivity of TRPA1 (48). The authors identified V875 in primate TRPA1, corresponding to G878 in rodents, and demonstrated that mutation G878V abolishes cold activation of the rat and mouse TRPA1. The role of this residue in cold-dependent gating was further supported by the analysis of the kinetic properties of the mutants of human TRPA1-V875G and mouse TRPA1-G878V at 12 °C, 25 °C and 35 °C (18). The other two cysteines, C1021 and C1025, are positioned within a short helix at the cytosol-membrane interface located near the S1 and S4 of the VSLD (43). A role for this structural motif (termed the interfacial helix, IFH) has been suggested in voltage-dependent gating and phospholipid regulation (18, 49). In the structure of TRPA1 captured in complex with the reversible covalent agonist benzyl isothiocyanate (PDB ID: 6PQP), the backbone carbonyl of C1025 directly interacts with N855, a residue preceding C856 and the site of a gain-of-function disease mutation causing syndrome characterized by pain that is triggered by physical stress, including noxious cold (34). Interestingly, the double mutant C1021A/C1025A restored hTRPA1 cold responses, and thus it is possible that the formation of a disulphide bond between C1021 and C1025 has a key impact on the C1025 – N855 interaction and channel gating. Together, this suggests the involvement of all the three cysteines 856, 1021, and 1025 in the agonist-, voltage- and cold-dependent gating of TRPA1. This further supports our findings that the redox state of hTRPA1 is important in determining its temperature-sensitive properties.

Two recent studies using cryo-EM as the core technique to explain thermoTRP gating did neither support nor dismissed the molar heat capacity model or an allosteric coupling model determined by specific amino acids by which TRPV1 and TRPV3 are gated by heat (10, 50). As proposed for TRPV1, both large global conformational changes of multiple subdomains followed by small amino acid specific outer pore conformational changes in a stepwise action are responsible for heat-dependent TRPV1 gate opening when capsaicin is bound in the vanilloid binding pocket that is formed by residues in the S2–S3 loop, S3, S4 and S4–S5 linker of one subunit, and S5 and S6 of the neighboring subunit (50). Likewise, heat-induced opening of TRPV3 is caused by a conformational wave involving the S2-S3 linker and the N and C-termini, and as a result modulation of the lipid occupying vanilloid binding site (10). Lipid displacement from the vanilloid pocket has also been proposed to cause heat activation of TRPV1 (51), and cholesterol binding to mouse TRPA1 influences its functional response to AITC (52). Interestingly, replacement with a single human amino acid in the pore region of drosophila TRPA1 reversed its thermosensitivity from heat to cold (53), and several VSLD lipid-binding pockets, also likely to bind carvacrol, are in close connection with CTD and N-ARD/pre-S1 region of hTRPA1 (31). Thus, in the case of hTRPA1, it will be interesting to further explore by cryo-EM if specific amino acids in the pore region are involved in cold-induced structural rearrangement, and if any of the lipid binding pockets in the VSLD is key for its heat activation. Naturally, the effect of redox on temperature-driven conformational changes and gating of hTRPA1 will also be of importance to study combining cryo-EM and mass spectrometry.

Our observation that point mutations at C856, C1021 and C1025 affect both voltage- and cold-sensitivity may fit with an allosteric model in which there is only one heat-activated temperature sensor (15). This model is based on the presumption that independent heat- and voltage-sensors exist that are coupled to the channel gate and to each other. The heat sensor is coupled to channel gating by a temperature-dependent allosteric coupling factor whose enthalpic contribution solely determines whether the channel is activated by both cooling and heating or only by cooling (Fig. 5 in (15)). We demonstrate that the deletion of the VSLD (Δ1-854 hTRPA1) led to a selective loss of the heat-activated branch of the relationship between the single-channel open probability and temperature. This could support the allosteric coupling model over the heat capacity model (14), as the latter predicts that the steepness of the cold- and heat-activated branches of the single-channel open probability - temperature relationship is symmetric. Even though a single temperature sensor e.g., located within the CTD would trigger both cold and heat responses, separate cold and heat pathways to the channel gate must exist as both functional bilayer experiments and tryptophan fluorescence studies of the purified hTRPA1s clearly showed that the VSLD is only needed for the heat sensitivity. However, it could also be that separate and specialized temperature sensor domains in the CTD, the conformational transformation of which may involve protein folding-unfolding, dictate the allosteric coupling of CTD to the gate (**Fig. 7**). A similar role for CTD has been proposed in the cold activation of TRPM8 (16), and may very well be involved in the temperature gating of TRPV1 and TRPV3 (10, 12, 50).

**Fig. 7.**
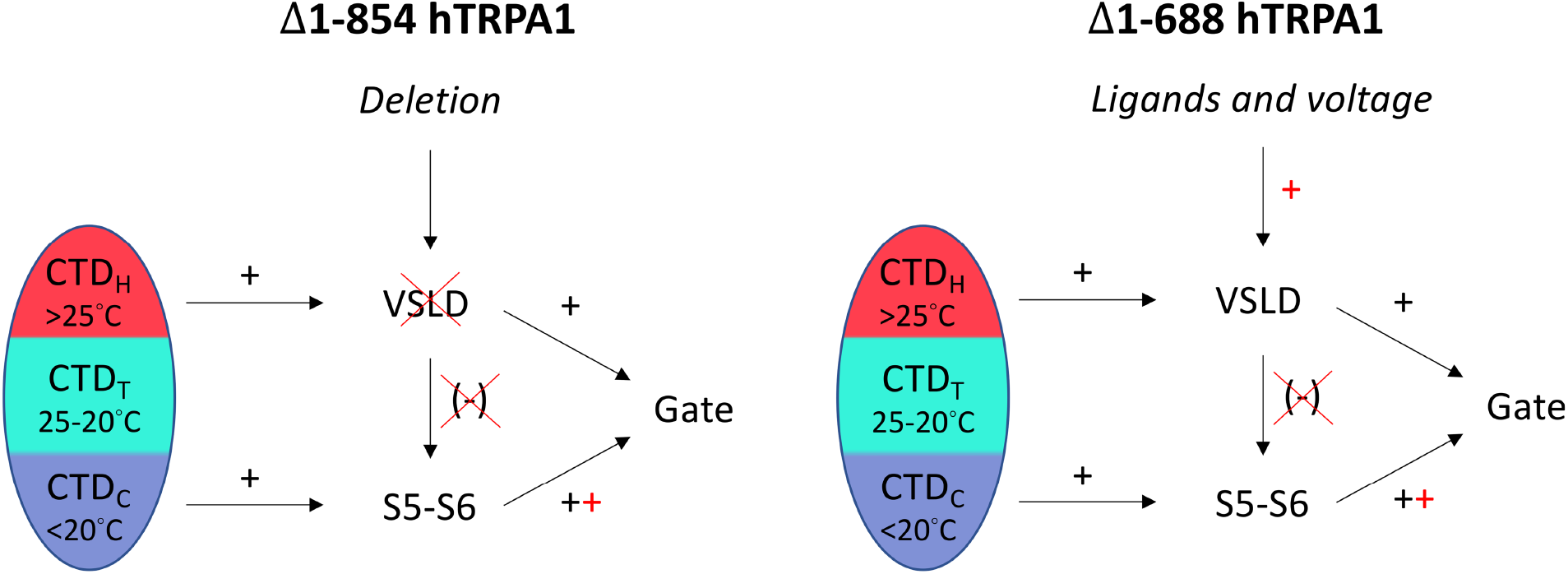
It is suggested that the hTRPA1 intracellular C-terminal domain (CTD) contains cold (CTD_C_) and heat (CTD_H_) sensitive domains, of which the CTD_C_ and S5-S6 transform cold exposure into channel opening whereas CTD_H_ together with the voltage-sensing like domain (VSLD) are involved in heat responses. The transition state (CTD_T_), under which CTD undergoes sudden major structural rearrangement, as could only be detected in the tryptophan fluorescence spectroscopy measurements of Δ1-854 hTRPA1 (containing only 2 tryptophans that are located within CTD), the temperature sensitivity of the channel is switched between heat and cold. When entering into the CTD_C_ state, further cooling opens the channel fully within a narrow temperature range of 5 to 10°C, but with minor changes in Δ1-854 hTRPA1 tryptophan fluorescence, suggesting that the CTD is in a stabilized state and conformational changes of S5-S6 occurs to open the gate. The same scenario seems valid for Δ1-688 hTRPA1 cold sensitivity, which also responded to cold with minor changes in tryptophan fluorescence in spite of additional 4 tryptophans of which 3 are within S3-S4 and 1 in pre-S1 (Fig. 4). In contrast to cold, heat caused substantial Δ1-854 hTRPA1 CTD structural rearrangements at temperatures above 25°C but without any increase in channel open probability, indicating that its heat sensitivity is not properly transmitted to the gate possibly by uncoupling of CTD_H_ and VSLD. Indeed, maintained heat sensitivity is associated with the complete quenching of the tryptophan fluorescence in Δ1-688 hTRPA1 indicating a major structural rearrangement of S1-S4, and supports that coupling between CTD_H_ and VSLD is crucial for proper heat activation of hTRPA1. The higher cold sensitivity of Δ1-854 hTRPA1 (Q_10_ = 1000) compared to Δ1-688 hTRPA1 (Q_10_ = 56) suggests that the VSLD restricts cold-induced channel conformations. Interestingly, the non-electrophilic TRPA1 activator carvacrol and voltage may remove this inhibitory effect by interacting with the VSLD. This may also be the case for oxidants and electrophilic compounds such as AITC, acrolein and NMM. Thus, the temperature sensitivity of TRPA1 and other thermoTRPs may be highly dependent on the cellular environment including channel redox state and other post-translational channel modifications.

The bidirectional temperature sensitivity of hTRPA1 may be a general property of all intrinsic wild-type thermoTRPs, as proposed by Clapham and Miller (14), but remains to be demonstrated. In this context, we did not find any activity of lipid bilayer reconstituted purified AgTRPA1(A) when lowering the temperature to 15°C (8). However, AgTRPA1(A) displayed basal activity even at 15°C when expressed in a cellular system (54). In comparison with AgTRPA1(B), AgTRPA1(A) has additional amino acids at the distal end of the N terminus reducing its heat sensitivity (54), and perhaps any cold sensitivity. Thus, it is possible that post-translational modifications as well as lower temperatures are needed to reveal any intrinsic AgTRPA1 cold sensitivity. Although TRPA1 stands out among thermoTRPs with regard to the many cysteines (28 in hTRPA1) and extreme sensitivity to electrophiles and oxidants (36, 37), other thermoTRPs such as TRPV1, TRPV2 and TRPM2 are redox sensitive with the ability to discriminate between various oxidants (22, 55-58). Thus, post-translational modifications including channel redox state are important to consider when the intrinsic functional and structural thermosensitive properties of TRP channels are studied to understand their role as true thermoTRPs in normal physiology and pathophysiology.

It is proposed that the CTD contains a bidirectional temperature switch priming hTRPA1 for either cold or heat, and that cold and heat responses are mediated by allosteric interactions between CTD and the S5-S6 pore region or the VSLD, respectively.

## Acknowledgements

This study was supported by the Swedish Research Council (2014-3801) and the Medical Faculty of Lund University – ALF (Dnr. ALFSKANE-451751), the Czech Science Foundation, (grant number 19-03777S).

## Author contributions

L.M. and P.M.Z. conceptualized and directed research; L.M., P.M.Z., M.F., V.V. designed research; L.M., V.S., M.F. performed research; L.M., V.S., M.F., V.V., P.M.Z. analyzed data; P.M.Z. drafted the paper and L.M., V.V. and P.M.Z. wrote the paper.

